# Phylogenomic analysis of the genus *Pseudomonas* and reclassification of *P. humi, P. zeshuii, P. psychrotolerans, P. nitritireducens, P. pharmacofabricae* and *P. panacis* are later heterotypic synonym of *P. citronellolis* Lang 2007, *P. luteola, P. oryzihabitans, P. nitroreducens* Lang 2007, *P. fluvialis* and *P. marginalis* (Brown 1918) Stevens 1925 (Approved Lists 1980), respectively

**DOI:** 10.1101/2021.05.19.444773

**Authors:** Ritu Rani Archana Kujur, Sushanta Deb, Subrata K Das

**Author notes:** **Correspondence:** Dr. Subrata K. Das, Institute of Life Sciences, Department of Biotechnology, Nalco Square, Bhubaneswar – 751023, India, /.

## Abstract

The present study described the comparative genomic analysis of the validly named species of the genus *Pseudomonas* to define the taxonomic assignment. Genomic information for 208 type strains was available in the NCBI genome database at the time of conducting this analysis. The ANI, AAI and *in silico* DNA DNA hybridization (isDDH) data were higher than the threshold values for the twelve strains with their closely related type species. Whole genome comparisons shared 97 - 99 % average nucleotide identity, 97.85 to 99.19 % average amino acid identity and 72.80 to 90.40 % digital DNA DNA hybridization values. Further, the phylogenomic analysis based on the core genome confirmed that *P. humi* CCA1 and *P. citronellolis* LMG 18378, *P. zeshuii* KACC 15471 and *P. luteola* NBRC 103146, *P. oryzihabitans* DSM 6835 and *P. psychrotolerans* DSM 15758, *P. nitroreducens* DSM 14399 and *P. nitritireducens* WZBFD3-5A2, *P. fluvialis* CCM 8778 and *P. pharmacofabricae* ZYSR67-Z, *P. panacis* DSM 18529 and *P. marginalis* DSM 13124 formed a monophyletic clade. Thus, we proposed six type species viz., *P. humi* CCA1, *P. zeshuii* KACC 15471, *P. psychrotolerans* DSM 15758, *P. nitritireducens* WZBFD3 5A2, *P. pharmacofabricae* ZYSR67 Z and *P. panacis* DSM 18529 are the later heterotypic synonym of *P. citronellolis* Lang 2007, *P. luteola, P. oryzihabitans, P. nitroreducens* Lang 2007, *P. fluvialis* and *P. marginalis* (Brown 1918) Stevens 1925 (Approved Lists 1980), respectively considering the priority date of publication.

---

*Pseudomonas* is a genus of Gram-negative, Gammaproteobacteria, belonging to the family Pseudomonadaceae and containing 246 validly described species as on 5 March 2021 were taxonomically characterized (https://lpsn.dsmz.de/genus/pseudomonas). Taxonomic assignment of the species primarily done using the polyphasic approach comprising 16S rRNA gene sequence similarity, phenotypic and chemotaxonomic analyses. The members of the genus *Pseudomonas* demonstrate a great deal of metabolic diversity and are able to colonize a wide range of niches. Recently, 16S rRNA sequence analysis has redefined the taxonomy of many bacterial species [1]. As a result, the genus *Pseudomonas* includes strains formerly classified in the genera *Chryseomonas* and *Flavimonas* [2]. Moreover, phylogeny using core genes protein coding sequences along with evaluation of average nucleotide identity (ANI) and in silico DDH are being used for the reclassification of several bacterial taxa [3,4]. It had been possible by whole genome sequencing of microorganisms and their comparative phylogenomic analysis. In this study, we carried out the phylogenomic analysis using the available whole genome sequence data of 208 validly named *Pseudomonas* species to check their taxonomic assignments.

Out of the 246 validly named type strains, genomic information for 208 *Pseudomonas* species was present in the NCBI genome database (last accessed March 5, 2021). 16S rRNA gene sequences of the validly named species of the genus *Pseudomonas* were retrieved from genome using barrnap program (https://github.com/tseemann/barrnap) and their similarity was calculated using pairwise nucleotide sequence alignment tool embedded in EzBioCloud server [5]. The 16S rRNA sequences were aligned using CLUSTAL W algorithm of MEGA7 software package [6]. Strains had average nucleotide identity (ANI) value more than or equal to 90% were considered for the construction of 16S rRNA phylogenetic tree. Evolutionary tree were calculated using maximum likelihood (ML) and maximum parsimony (MP) algorithm by the Kimura two-parameter [7] and the topologies of the resultant trees were evaluated by bootstrap analyses [8] based on 1000 replicates. 16S rRNA gene sequence similarity between *Pseudomonas humi* CCA1 vs *Pseudomonas citronellolis* LMG 18378 was (99.45%), between *Pseudomonas zeshuii* KACC 15471 vs *Pseudomonas luteola* NBRC 103146 (99.8%), between *Pseudomonas oryzihabitans* DSM 6835 vs *Pseudomonas psychrotolerans* DSM 15758 (100%), between *Pseudomonas nitroreducens* DSM 14399 vs *Pseudomonas nitritireducens* WZBFD3-5A2 (99.87%), between *Pseudomonas fluvialis* CCM 8778 vs *Pseudomonas pharmacofabricae* ZYSR67-Z (99.87%) and between *Pseudomonas panacis* DSM 18529 vs *Pseudomonas marginalis* DSM 13124 (99.87%), respectively. The 16S rRNA gene sequence obtained from the genome of all the compared strains shared more than 99% similarity, which was higher than the recently corrected threshold value (98.65%) to define a bacterial species [9]. In the phylogenetic tree, strains that are showing 16S rRNA gene sequence similarity more than 99% were clustered together in the same clade. The maximum likelihood phylogenetic tree presented in **Fig. 1** is similar to the dendrogram generated by maximum-parsimony algorithms (**Fig. S1**) indicated robustness of tree topologies.

**Fig 1.**
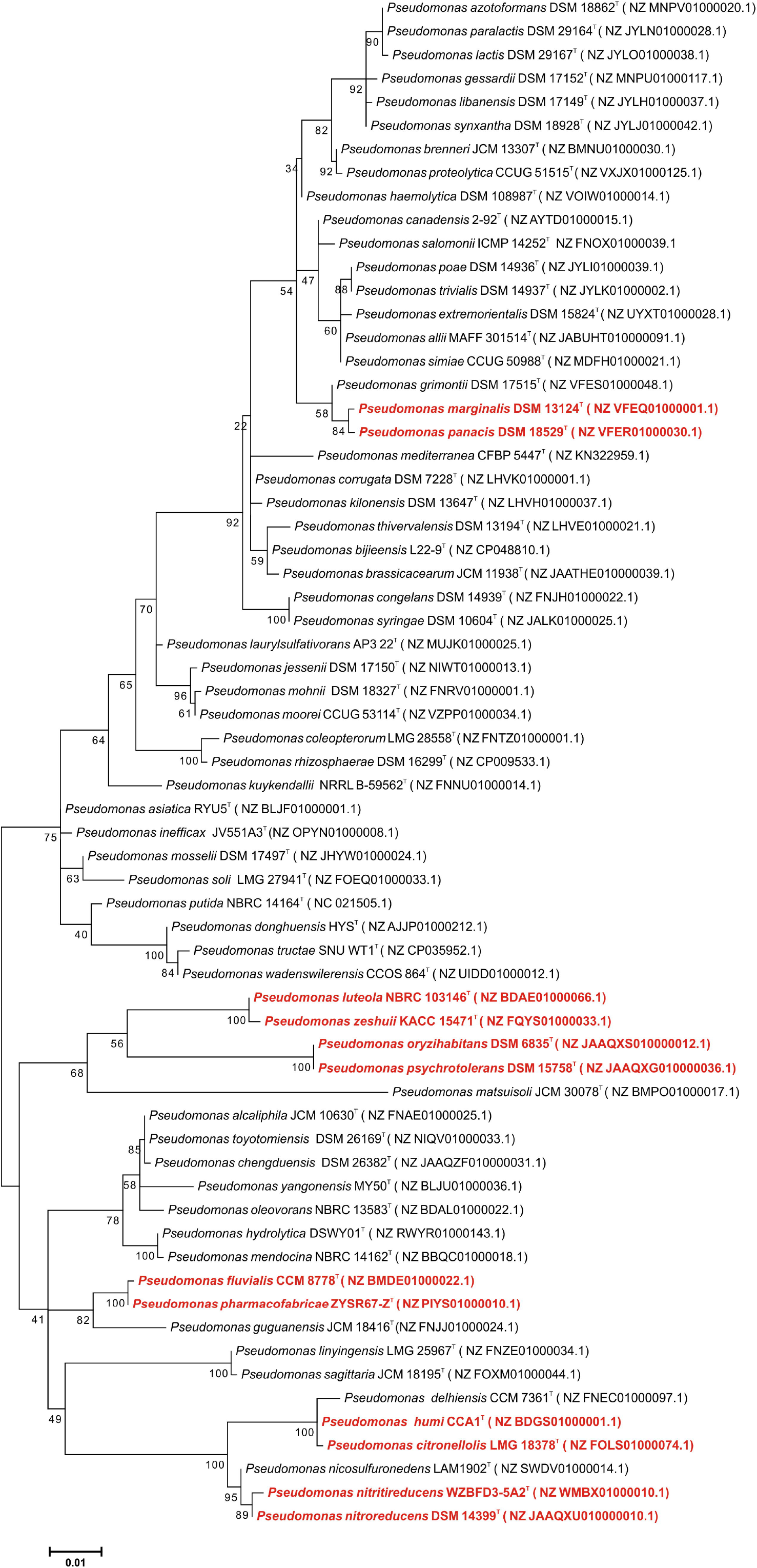
Maximum likelihood phylogenetic tree based on 16S rRNA gene sequence showing the heterotypic synonym strains are clustered in a monophyletic clade. Bootstrap values expressed as percentages of 1000 replications are given at branching points. Accession numbers are given in parentheses. Bar 1 substitution per 100 nucleotide positions.

Further, we performed genome-based comparison. Several bioinformatics analysis tools were used to compare the genomic relatedness of the validly named species of the genus *Pseudomonas* available in the National Centre for Biotechnology Information (NCBI) database (last accessed March 5, 2021). Comparison of the whole genome sequence characteristics of the reference strains compared in this study and proposed as a later heterotypic synonyms are listed in **Table-1**. The use of whole genome sequences has been regarded as a promising avenue for the taxonomic and phylogenetic studies of microorganisms. With the advent of next-generation sequencing and bioinformatics tools make possible to compare genomic data by *in silico* DDH (*is*DDH), average nucleotide identity (ANI) and average amino acid identity (AAI) values. Thus, we evaluated all the twelve validly named species based on the similarity in their genomic data.

**Table 1.** Genome statistics of the *Pseudomonas* strains compared in this study.

| SI. No. | Strains | Accession no. | Number of Scaffolds | GC (%) | Size (Mb) | Genes | Proteins | Completeness | Contamination |
| --- | --- | --- | --- | --- | --- | --- | --- | --- | --- |
| 1 | *Pseudomonas humi* CCA1 | BDGS000000000 | 24 | 67.2 | 6.99 | 6202 | 6029 | 100 | 3.63 |
| 2 | *Pseudomonas citronellolis* LMG 18378 | FOLS000000000 | 76 | 67.5 | 7.19 | 6435 | 6293 | 100 | 2.03 |
| 3 | *Pseudomonas luteola* NBRC 103146 | BDAE000000000 | 104 | 55.3 | 5.39 | 5059 | 4919 | 99.62 | 1.07 |
| 4 | *Pseudomonas zeshuii* KACC 15471 | FQYS000000000 | 33 | 55.3 | 5.33 | 4979 | 4800 | 99.62 | 1.08 |
| 5 | *Pseudomonas psychrotolerans* DSM 15758 | JAAQXG000000000 | 40 | 65.3 | 6.01 | 5710 | 5554 | 99.67 | 0.72 |
| 6 | *Pseudomonas oryzihabitans* DSM 6835 | JAAQXS000000000 | 13 | 66.2 | 5.03 | 4683 | 4574 | 100 | 0.11 |
| 7 | *Pseudomonas nitritireducens* WZBFD3-5A2 | WMBX000000000 | 12 | 64.8 | 6.39 | 5890 | 5776 | 100 | 1.73 |
| 8 | *Pseudomonas nitroreducens* DSM 14399 | JAAQXU000000000 | 50 | 65 | 6.17 | 5699 | 5595 | 99.98 | 1.78 |
| 9 | *Pseudomonas pharmacofabricae* ZYSR67-Z | PIYS000000000 | 34 | 62.6 | 3.43 | 3341 | 3212 | 96 | 0.14 |
| 10 | *Pseudomonas fluvialis* CCM 8778 | BMDE000000000 | 28 | 62.6 | 3.31 | 3142 | 3047 | 96 | 0.27 |
| 11 | *Pseudomonas marginalis* DSM 13124 | VFEQ000000000 | 81 | 60.6 | 7.18 | 6807 | 6537 | 99.93 | 1.15 |
| 12 | *Pseudomonas panacis* DSM 18529 | VFER000000000 | 46 | 61.1 | 6.80 | 6331 | 6155 | 99.93 | 0.34 |

The average nucleotide identity (ANI) was calculated using the python module pyani (https://github.com/widdowquinn/pyani) with ANIb method. In silico DDH similarity was measured with the help of Genome-to-Genome Distance calculator (formula 2) [10]. Average amino acid identity (AAI) was estimated using the ‘aai_wf’ function implemented in compareM program (https://github.com/dparks1134/CompareM). For phylogenomic analysis using core genomes of *Pseudomonas*, a comprehensive database of 208 whole genome sequences of different *Pseudomonas* species were retrieved from the NCBI database using the ncbi-genome-download tool (https://github.com/kblin/ncbi-genome-download/). The core genes extracted by UBCG pipeline [11] was concatenated and a maximum-likelihood tree reconstructed with the GTR model using RAxML tool [12]. Further, the non-recombinant core-genome based phylogenetic tree was constructed following the method of Mateo-Estrada *et al* [13]. The average nucleotide identity (ANI), average amino acid identity (AAI) and *in silico* DNA-DNA similarity values between *Pseudomonas humi* CCA1 vs *Pseudomonas citronellolis* LMG 18378, between *Pseudomonas zeshuii* KACC 15471 vs *Pseudomonas luteola* NBRC 103146, between *Pseudomonas oryzihabitans* DSM 6835 vs *Pseudomonas psychrotolerans* DSM 15758, between *Pseudomonas nitroreducens* DSM 14399 vs *Pseudomonas nitritireducens* WZBFD3-5A2, between *Pseudomonas fluvialis* CCM 8778 vs *Pseudomonas pharmacofabricae* ZYSR67-Z and between *Pseudomonas panacis* DSM 18529 vs *Pseudomonas marginalis* DSM 13124 are listed in **Table 2.** The calculated ANI (97.0 to 99.0%) values between pair of strains compared in the present study are higher than the proposed thresholds values (95–96%) for bacterial species delineation **(supplementary dataset)** [14, 15]. Phylogenomic tree based on concatenated core genes (**Fig. 2**) showed that *P. humi* CCA1 and *P. citronellolis* LMG 18378, *P. zeshuii* KACC 15471 and *P. luteola* NBRC 103146, *P. oryzihabitans* DSM 6835 and *P. psychrotolerans* DSM 15758, *P. nitroreducens* DSM 14399 and *P. nitritireducens* WZBFD3-5A2, *P. fluvialis* CCM 8778 and *P. pharmacofabricae* ZYSR67-Z, *P. panacis* DSM 18529 and *P. marginalis* DSM 13124 clustered together. Moreover, the non-recombinant core-genome based phylogenetic tree revealed these strains were clustered in the same clade (**Fig. 3)** compared to the tree generated using core genomes to determine the taxonomic position (**Fig. 2**), indicating robustness of tree topology. Further, in silico DDH value between the strains was higher than the 70% cut off to define bacterial species [10]. Moreover, the average amino acid identity (AAI) between the strains were more than 97.85% **(Table 2).** These analysis suggest that the two type strains *P. humi* CCA1 and *P. citronellolis* LMG 18378, *P. zeshuii* KACC 15471 and *P. luteola* NBRC 103146, *P. oryzihabitans* DSM 6835 and *P. psychrotolerans* DSM 15758, *P. nitroreducens* DSM 14399 and *P. nitritireducens* WZBFD3-5A2, *P. fluvialis* CCM 8778 and *P. pharmacofabricae* ZYSR67-Z, *P. panacis* DSM 18529 and *P. marginalis* DSM 13124, respectively are actually belong to the same species. *P. citronellolis* Lang *et al* [16] have priority over the names *Pseudomonas humi* [17]. Similarly, *Pseudomonas luteola* Kodama *et al* [18] have priority over the names *Pseudomonas zeshuii* [19], *Pseudomonas oryzihabitans* Kodama *et al* [18] have priority over the names *Pseudomonas psychrotolerans* [20], *Pseudomonas nitroreducens* Lang *et al* [16] have priority over the names *Pseudomonas nitritireducens* [23], *Pseudomonas fluvialis* Sudan *et al* [25] have priority over the names *Pseudomonas pharmacofabricae* [24] and *Pseudomonas marginalis* (Brown 1918) Stevens 1925 (Approved Lists 1980) [26] have priority over the names *Pseudomonas panacis* [27].

**Fig 2.**
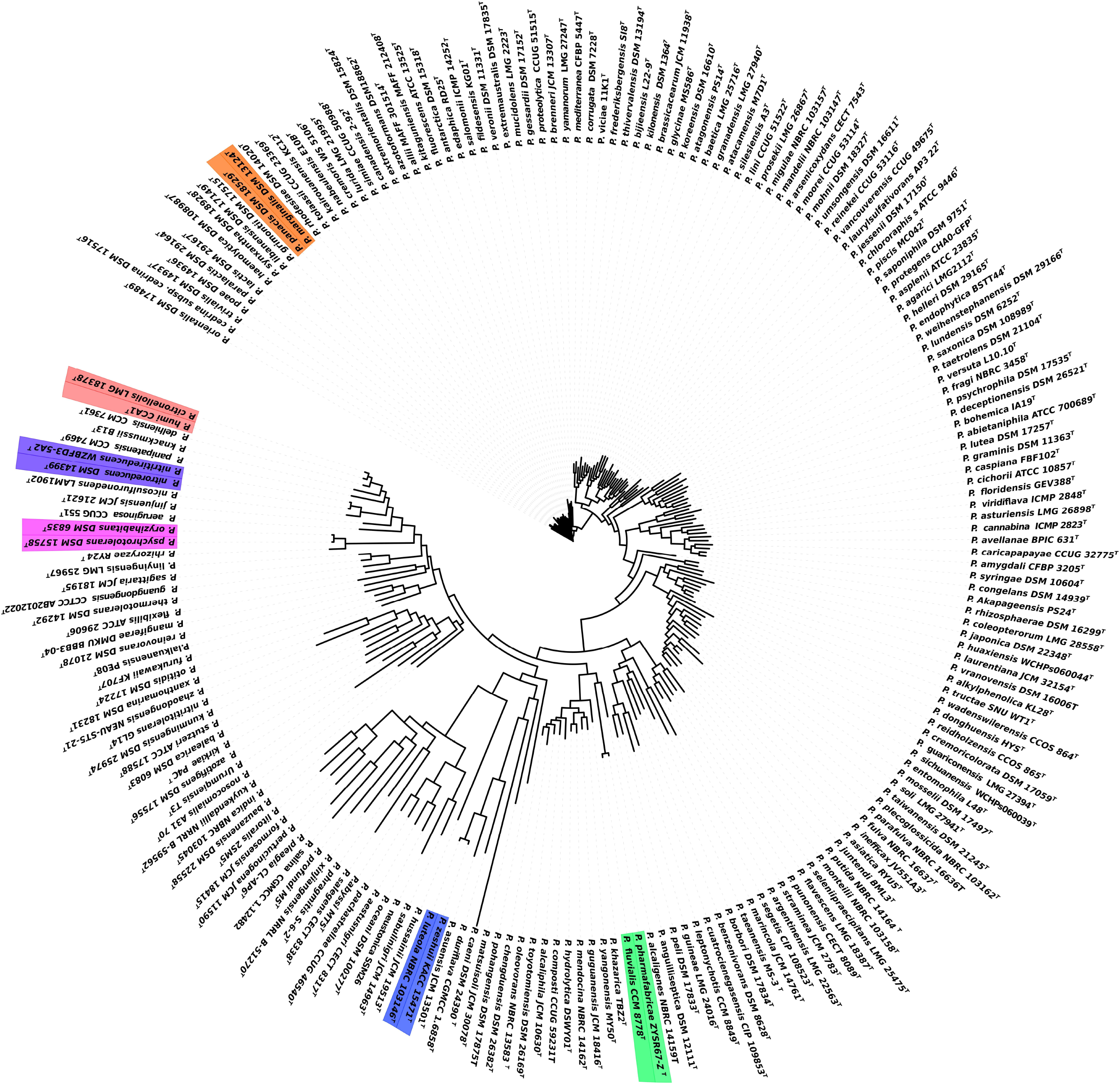
Phylogenetic tree based on the concatenated core genes from 208 type strains of *Pseudomonas*. Phylogenetic position of the heterotypic synonym strains are clustered in a monophyletic clade.

**Fig 3.**
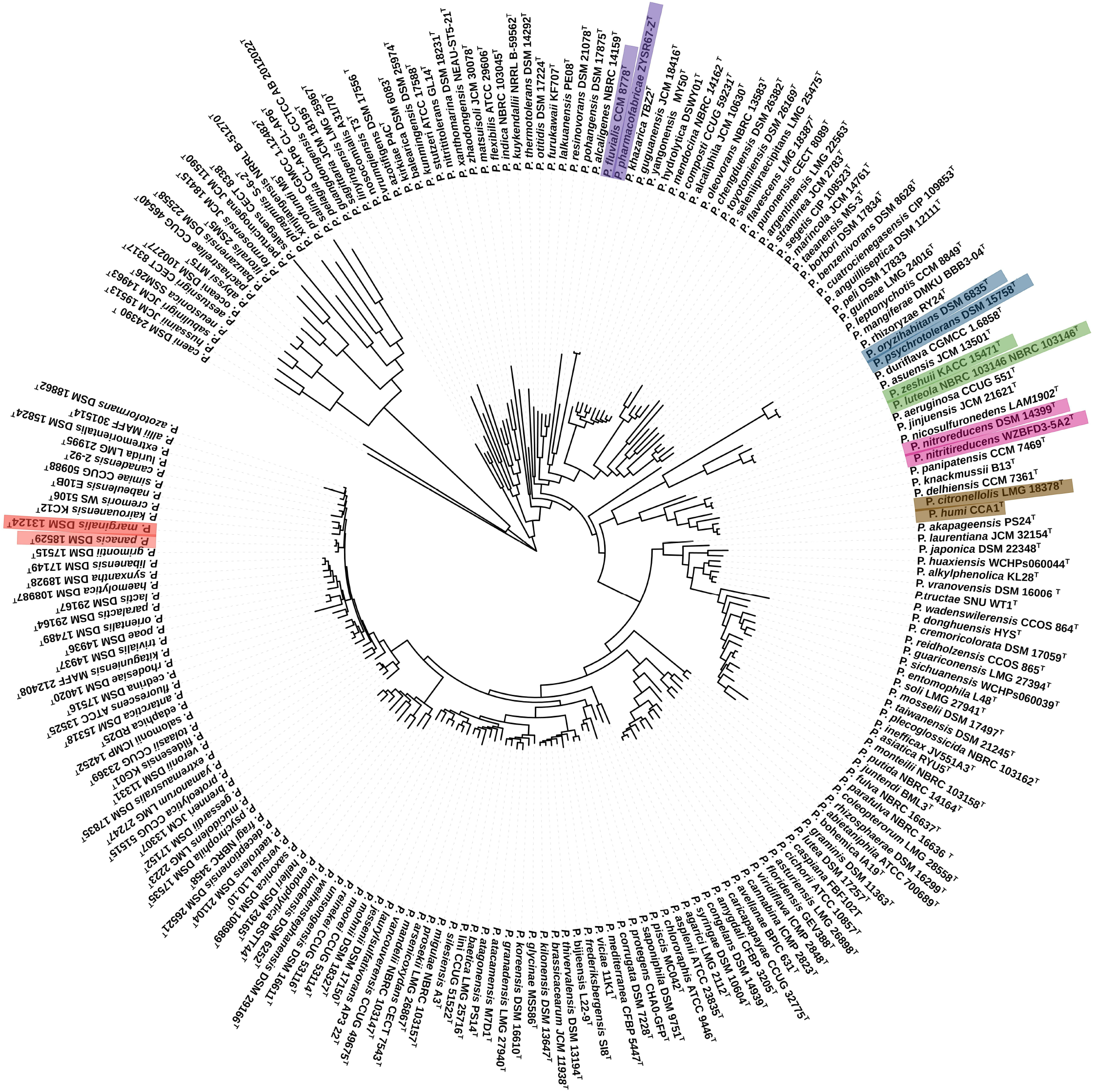
Non-recombinant core-genome based phylogenetic tree from 208 type strains of *Pseudomonas*. Phylogenetic position of the heterotypic synonym strains are clustered in a monophyletic clade.

**Table 2.** Overall genomic indices between heterotypic synonym strains of *Pseudomonas* clustered together in the phylogenetic tree.

| SI. no | *Pseudomonas* strains | ANI | AAI | DDH (%) similarity | 16S rRNA (%) similarity |
| --- | --- | --- | --- | --- | --- |
| 1 | *Pseudomonas humi* CCA1<br>vs<br>*Pseudomonas citronellolis* LMG 18378 | 97 | 97.85 | 72.80 | 99.45 |
| 2 | *Pseudomonas luteola* NBRC 103146<br>vs<br>*Pseudomonas zeshuii* KACC 15471 | 98 | 98.51 | 82 | 99.8 |
| 3 | *Pseudomonas psychrotolerans* DSM 15758<br>vs<br>*Pseudomonas oryzihabitans* DSM 6835 | 98 | 98.59 | 86.10 | 100 |
| 4 | *Pseudomonas nitritireducens* WZBFD3-5A2<br>vs<br>*Pseudomonas nitroreducens* DSM 14399 | 99 | 99.19 | 90.40 | 99.87 |
| 5 | *Pseudomonas pharmacofabricae* ZYSR67-Z<br>vs<br>*Pseudomonas fluvialis* CCM 8778 | 99 | 98.89 | 88.20 | 99.87 |
| 6 | *Pseudomonas marginalis* DSM 13124<br>vs<br>*Pseudomonas panacis* DSM 18529 | 97 | 98.00 | 74.90 | 99.87 |

Based on the genomic analysis, we therefore propose that *P. humi, P. zeshuii, P. psychrotolerans, P. nitritireducens, P. pharmacofabricae* and *P. panacis* are later heterotypic synonym of *P. citronellolis* Lang 2007, *P. luteola, P. oryzihabitans, P. nitroreducens* Lang 2007, *P. fluvialis* and *P. marginalis* (Brown 1918) Stevens 1925 (Approved Lists 1980), respectively.

## EMENDED DESCRIPTION OF *PSEUDOMONAS CITRONELLOLIS*

The description is as before [16] with the following modification. *Pseudomonas humi* Akita *et al*. 2019 is a later heterotypic synonyms of *Pseudomonas citronellolis* Lang *et al*. 2007.

Type strain is ATCC 13674^T^ (= CCUG 17933^T^ = CFBP 5585^T^ = CIP 104381^T^ = DSM 50332^T^ = IAM 15129^T^ = JCM 21587^T^ = LMG 18378^T^ = NBRC 103043^T^ = NRRL B-2504^T^).

## EMENDED DESCRIPTION OF *PSEUDOMONAS LUTEOLA*

The description is as before [18] with the following modification. *Pseudomonas zeshuii* Feng *et al*. 2012 is a later heterotypic synonyms of *Pseudomonas luteola* Kodama *et al*. 1985.

Type strain is 4239^T^ (= ATCC 43273^T^ = CCUG 37974^T^ = CFBP 3007^T^ = CIP 102995^T^ = DSM 6975^T^ = IAM 1300^T^ = IAM 13000^T^ = JCM 3352^T^ = LMG 7041^T^ = NBRC 103146^T^)

## EMENDED DESCRIPTION OF *PSEUDOMONAS ORYZIHABITANS*

The description is as before [18] with the following modification. *Pseudomonas psychrotolerans* Hauser *et al*. 2004 is a later heterotypic synonyms of *Pseudomonas oryzihabitans* Kodama *et al*. 1985.

Type strain is AJ 2197^T^ (= ATCC 43272^T^ = CCUG 12540^T^ = CIP 102996^T^ = DSM 6835^T^ = IAM 1568^T^ = JCM 2952^T^ = LMG 7040^T^ = NBRC 102199^T^).

## EMENDED DESCRIPTION OF *PSEUDOMONAS NITROREDUCENS*

The description is as before [16] with the following modification. *Pseudomonas nitritireducens* Wang *et al*. 2012 is a later heterotypic synonyms of *Pseudomonas nitroreducens* Lang *et al*. 2007.

Type strain is WZBFD3-5A2^T^ (= CGMCC 1.10702^T^ = LMG 25966^T^).

## EMENDED DESCRIPTION OF *PSEUDOMONAS FLUVIALIS*

The description is as before [25] with the following modification. *Pseudomonas pharmacofabricae* Yu *et al*. 2018 is a later heterotypic synonyms of *Pseudomonas fluvialis* Sudan et *al*. 2018

Type strain is ASS-1^T^ (= CCM 8778^T^ = KCTC 52437^T^).

## EMENDED DESCRIPTION OF *PSEUDOMONAS MARGINALIS*

The description is as before [26] with the following modification. *Pseudomonas panacis* (Park *et al*. 2005) is a later heterotypic synonyms of *Pseudomonas marginalis* (Brown 1918) Stevens 1925 (Approved Lists 1980).

The type strain is ATCC 10844^T^ (= CFBP 1387^T^ = CFBP 2037^T^ = CIP 106712^T^ = DSM 13124^T^ = ICMP 3553^T^ = LMG 2215^T^ = NCPPB 667^T^).

## Acknowledgments

The author RRAK and SD acknowledges the Council of Scientific and Industrial Research (CSIR), New Delhi, Government of India and Institute of Life Sciences, Bhubaneswar for providing the research fellowship. We acknowledge the Distributed Information Sub-Center (DISC) at Institute of Life Sciences, Bhubaneswar for the computational facility.

## Authors and contributors

SKD developed the concept, RRAK and SD conducted the experiment. RRAK, SD and SKD analysed the data and wrote the manuscript.

## Ethical approval

Ethical approval not required for this study

