## Supplemental figure for "Phylogenomic analysis of the genus *Pseudomonas* and reclassification of *P. humi, P. zeshuii, P. psychrotolerans, P. nitritireducens, P. pharmacofabricae* and *P. panacis* are later heterotypic synonym of *P. citronellolis* Lang 2007, *P. luteola, P. oryzihabitans, P. nitroreducens* Lang 2007, *P."

***Correspondence:**

Dr. Subrata K. Das

Phone: (+91) 6742304328, Fax: (+91) 6742300728

**Supplementary Figure Legends**

**Fig. S1.**	Maximum parsimony phylogenetic tree based on 16S rRNA gene sequence showing the heterotypic synonym strains are clustered in a monophyletic clade.

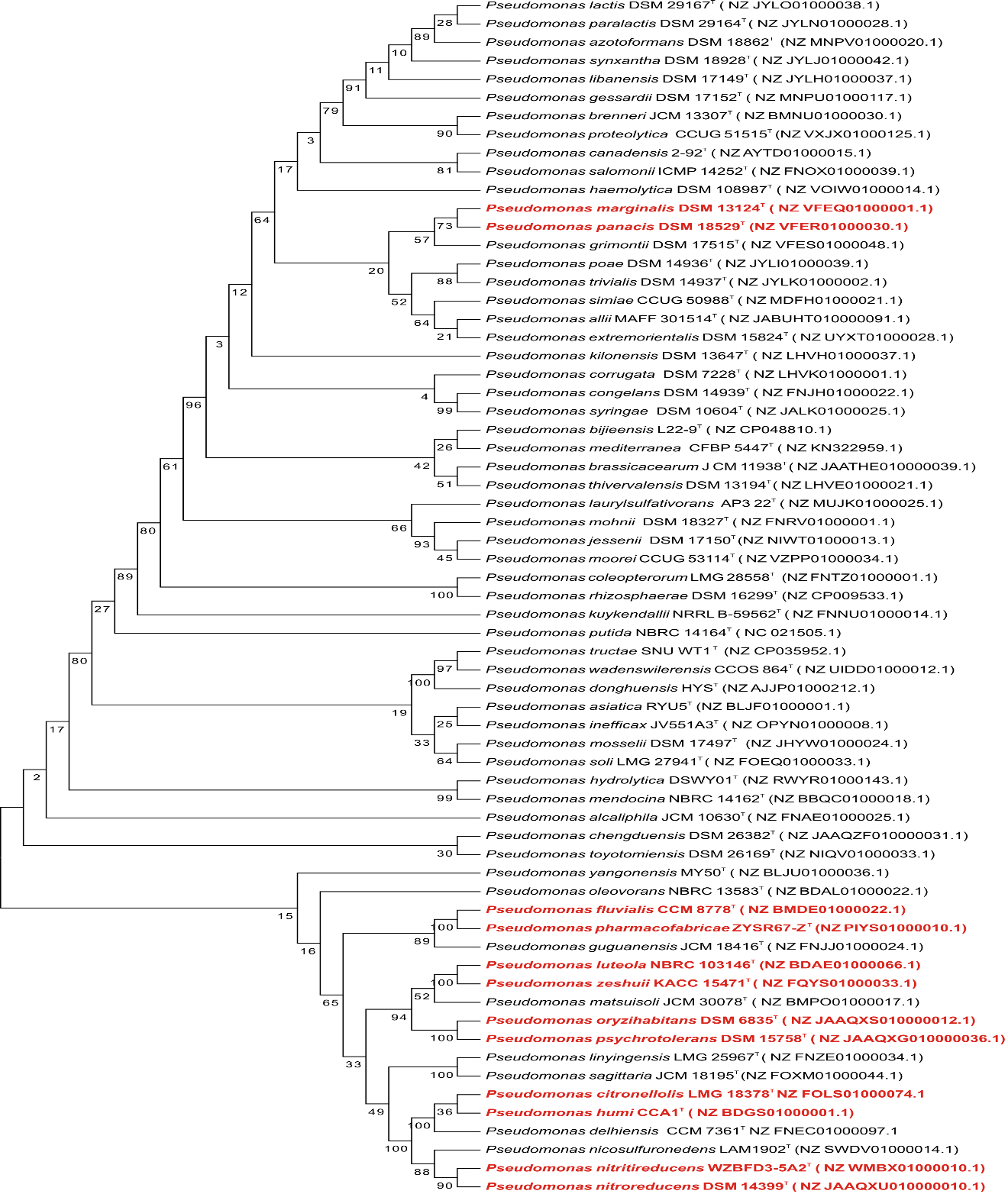

**Fig. S1.**
